# LCM-seq reveals unique transcriptional adaption mechanisms of resistant neurons in spinal muscular atrophy

**DOI:** 10.1101/356113

**Authors:** S Nichterwitz, H Storvall, J Nijssen, LH Comley, I Allodi, M van der Lee, C Schweingruber, Q Deng, R Sandberg, E Hedlund

## Abstract

Somatic motor neurons are selectively vulnerable in spinal muscular atrophy (SMA), a lethal disease caused by a deficiency of the ubiquitously expressed survival of motor neuron (SMN) protein. However, some brainstem motor neuron groups, including oculomotor and trochlear (ocular), which innervate the muscles around the eyes, are for unknown reasons spared. Here, using laser capture microdissection coupled with RNA sequencing (LCM-seq), we investigate the transcriptional dynamics in discrete neuronal populations in health and SMA to reveal mechanisms of vulnerability and resistance. Using gene correlation network analysis, we reveal a p53-mediated stress response that is intrinsic to all somatic motor neurons independent of their vulnerability, but absent in resistant red nucleus and visceral motor neurons. However, our temporal and spatial differential expression analysis across neuron types clearly demonstrates that the majority of SMA-induced modulations are cell-type specific. Notably, using gene ontology and protein-network analyses we show that ocular motor neurons present unique disease-adaptation mechanisms that could explain their resilience. In particular, ocular motor neurons up-regulate; *i*) *Syt1*, *Syt5* and *Cplx2*, which modulate neurotransmitter release; *ii*) the motor neuron survival factors *Chl1 and Lif, iii) Aldh4*, that can protect cells from oxidative stress and *iv*) the caspase inhibitor *Pak4*. In conclusion, our in-depth longitudinal analysis of gene expression changes in SMA reveal novel cell-type specific changes that present compelling targets for future gene therapy studies aimed towards preserving vulnerable motor neurons.

## Introduction

Spinal muscular atrophy (SMA) is an autosomal recessive disease, characterized by the progressive degeneration of somatic motor neurons in spinal cord and lower brainstem. SMA is caused by the loss of functional survival motor neuron (SMN) protein due to loss of or mutations in the telomeric gene *SMN1*. SMA displays a wide clinical spectrum and is classified based on age of onset and severity of the disease. An increased copy number of the centromeric *SMN2* gene is the main predictor of disease severity (Feldkötter et al. 2002; Vitali et al. 1999; Lorson et al. 1999; Wirth 2000). *SMN1* and *SMN2* differ by five nucleotides only and encode identical proteins. However, the C to T nucleotide transition in exon 7 of the *SMN2* gene disrupts an exonic splicing enhancer and leads to alternative splicing, in which a majority of *SMN2* transcripts lack exon 7 (*SMN*Δ*7*) (Monani et al. 1999; Lorson et al. 1999). While SMNΔ7 appears to be a functional SMN protein, it is highly unstable and quickly degraded (Cho and Dreyfuss 2010). Recently, the first drug treatment for SMA, based on increasing full-length SMN from *SMN2* exon 7 inclusion, was approved. This presents a very promising therapeutic strategy for the broad treatment of SMA, with positive outcomes in several clinical studies (Finkel et al. 2017; Pechmann et al. 2018). However, the timing of the initiation of treatment appears crucial for the outcome and patients will very likely benefit from additional treatments that aim to preserve or improve motor function.

The best characterized function of SMN is its role in the assembly of small nuclear ribonucleoproteins (snRNPs), which are major components of the pre-mRNA splicing machinery (Fischer et al. 1997). SMN can be found in nuclear gems and in the cytoplasm, but in neurons it is also located in axons and growth cones (Pagliardini et al. 2000; Giavazzi et al. 2006; Jablonka et al. 2001). A more widespread role for SMN in RNP assembly is now accepted due to the disruption of axonal mRNA localization and translation in an SMN-deficient context (Donlin-Asp et al. 2016). There is also strong evidence that SMN can prevent DNA damage and apoptosis (Vyas et al. 2002; Zhao et al. 2015; Sareen et al. 2012). As of now, it remains unclear which of these functions, when disrupted, lead to SMA.

SMN is ubiquitously expressed and its complete depletion leads to death in early embryonic development (Schrank et al. 1997). In SMA, somatic motor neurons are for unknown reasons selectively vulnerable to the lower level of SMN protein. However, different motor neuron groups show varying degrees of susceptibility to degeneration. Spinal motor neurons are the primarily affected cell type in disease. Facial motor neurons and hypoglossal motor neurons that innervate the tongue are to some extent affected in severe cases of the human disease (Rudnik-Schonebörn et al. 2009; Petit et al. 2011; Harding et al. 2015). However, whereas neuromuscular junctions (NMJs), the specialized synapses between motor neurons and muscle, of facial motor neurons present pathology in mouse models of SMA (Murray et al. 2008), hypoglossal NMJs remain unaffected (Comley et al. 2016). Ocular motor neurons, which innervate extraocular muscles and thus control movement of the eyes, appear resistant in SMA. This is evidenced by the use of ocular tracking devices as a communication tool for patients (Kubota et al. 2000). Consistently, we have shown that NMJs are preserved in extraocular muscles of end-stage SMA mice (Comley et al. 2016).

Genes active within specific neuronal types define their unique identities and functions in health as well as their susceptibility to specific neurodegenerative diseases. Importantly, data from SMA animal models and SMA patient motor neurons derived from induced pluripotent stem cells (iPSCs), indicate that factors intrinsic to motor neurons are important for degeneration (Park et al. 2010; Corti et al. 2012; Van Hoecke et al. 2012). Thus, investigating cell intrinsic pathways that are differentially activated within resistant and vulnerable motor neurons could reveal mechanisms of selective neuronal degeneration and lead to therapies preventing progressive motor neuron loss (Nijssen et al. 2017; Allodi et al. 2016; Comley et al. 2015; Hedlund et al. 2010).

Previous transcriptome studies in SMA have compared vulnerable patient-derived motor neurons (Ng et al. 2015) or whole spinal cords isolated from SMA mice (Staropoli et al. 2015; Murray et al. 2010; Bäumer et al. 2009; Zhang et al. 2008) with healthy controls. These studies have improved our understanding of motor neuron disease mechanisms, but could not explain how the loss of a ubiquitously expressed protein can induce degeneration in a select cell type. Zhang et al. (2013) included unaffected cells from the white matter in their analysis and a more recent study by Murray et al. (2015) investigated transcriptional changes in differentially affected motor neuron pools. However, these studies were restricted to a single pre-symptomatic stage, limiting the scope of the findings. To unravel temporal mechanisms of neuronal resilience and susceptibility we conducted a comprehensive longitudinal analysis of resistant and vulnerable neuron groups from a pre-symptomatic stage to early and late symptomatic stages. We used laser capture microdissection coupled with RNA sequencing (LCM-seq) (Nichterwitz et al. 2016, 2018) to profile discrete neuronal populations in SMA mice and littermate controls over time. Our findings provide important insight into common and cell-type specific transcriptome changes. Our study highlights the importance of p53 signaling and the DNA damage response in the pathology of SMA, that appears intrinsic to somatic motor neurons. Importantly, we reveal transcriptional changes restricted to resistant oculomotor and trochlear motor neurons that could contribute to their resistance. Among the specifically upregulated genes were *Syt1*, *Syt5*, *Chl1*, *Cplx2*, *Aldh4* and *Pak4*, that present attractive targets for further investigation in the context of selective motor neuron susceptibility in SMA.

## Results

### Transcriptional profiling of neurons with differential susceptibility to degeneration reveals cell-type specific gene expression

To investigate the transcriptional dynamics of neuronal populations with differential vulnerability in SMA we used the widely studied ‘delta7’ mouse model (*Smn*^−/−^/*SMN2*^+/+^/*SMN*Δ*7*^+/+^). As we were interested in longitudinal changes in gene expression we analyzed several disease stages. We included a pre-symptomatic stage (P2) and an early symptomatic stage (P5), when motor neuron loss in this model is restricted to discrete regions of the spinal cord. We also included a symptomatic stage (P10) when the *Smn* deficient mice have clear motor dysfunction and show a more widespread loss of spinal motor neurons (Le et al. 2005; Mentis et al. 2011) (Fig. 1A). We applied laser capture microdissection (LCM) to isolate neurons from different regions of the brainstem and spinal cord (Fig. 1A, Supplemental Fig. S1) coupled with poly(A)-enriched RNA sequencing (LCM-seq; (Nichterwitz et al. 2016, 2018)). We collected somatic motor neurons from the oculomotor and trochlear nuclei (cranial nuclei 3 and 4 (CN3/4), Supplemental Fig. S1D-F) and the hypoglossal nucleus (cranial nucleus 12 (CN12), Supplemental Fig. S1M-O) that are resistant to degeneration in this SMA model. We also isolated vulnerable somatic motor neurons from the facial nucleus (cranial nucleus 7 (CN7), Supplemental Fig. S1G-I) and lumbar spinal cord (SC, Supplemental Fig. S1P-R).

**Figure 1.**
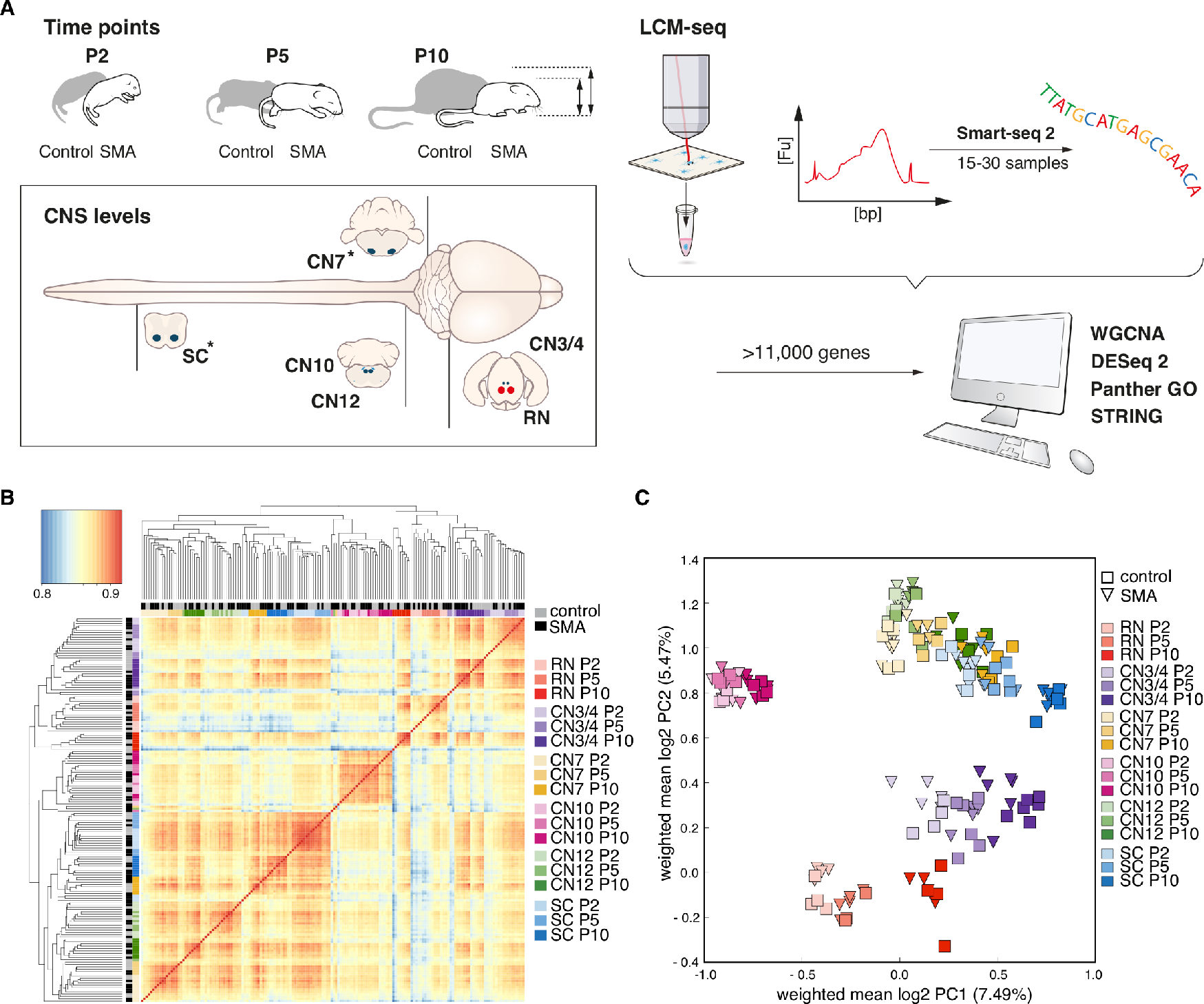
LCM-seq strategy to reveal cell intrinsic mechanisms of motor neuron vulnerability and resistance in SMA. *(A)* We used ‘delta 7’ mice (*Smn*^−/−^/*SMN2*^+/+^/*SMNΔ7*^+/+^) as a model for severe SMA and littermates homozygous for murine *Smn* as controls (*Smn*^+/+^/*SMN2*^+/+^/*SMNΔ7*^+/+^). *(B)* Pairwise Spearman correlation on log_10_-transformed data of all samples per cell type, genotype and age. *(C)* Principal component analysis on the whole gene expression data set. P = postnatal day; CNS = central nervous system; LCM-seq = laser capture microdissection coupled with RNA sequencing; SC = spinal cord, CN = cranial nerve; CN12 = hypoglossal nucleus; CN10 = dorsal motor nucleus of the vagus nerve; CN7 = facial nucleus; CN3/4 = oculomotor and trochlear nuclei; RN = red nucleus; asterisks in *A* indicate vulnerable cell types in this mouse model; WGCNA = weighted gene correlation network analysis; GO = gene ontology.

Furthermore, we collected resistant visceral motor neurons from the dorsal motor nucleus of the vagus nerve (vagus motor neurons) (cranial nucleus 10 (CN10), Supplemental Fig. S1J-L) to deduct events occurring within all cholinergic motor neurons versus those specific for somatic motor neurons. We also isolated red nucleus neurons (RN, Supplemental Fig. S1A-C), which are non-cholinergic neurons involved in motor coordination, to elucidate disease-induced transcriptional regulation selective to cholinergic neurons versus more broad regulation across neuronal populations. We thus acquired an extensive library of six neuronal populations at three different time points throughout disease progression in health and SMA.

After conducting LCM-seq, we performed a quality control, using only samples where we achieved a gene detection level of >11,000 expressed genes (Supplemental Fig. S2A), which left a total of 168 samples for further analysis (Supplemental Table S1). This control was done to exclude samples with lower RNA-seq library quality. To evaluate the purity of the LCM-seq samples, we analyzed the level of neuronal and glial markers and compared these to a previously published RNA sequencing data set of neurons, astrocytes, oligodendrocytes and microglia (GEO accession number GSE52564 (Zhang et al. 2014)). The neuronal markers neurofilament heavy chain (*Nefh*) and peripherin (*Prph*) were highly expressed in all our samples. The motor neuron markers choline acetyl transferase (*Chat*) and Islet-1/2 were readily detected in all motor neuron groups, while *Hb9* (*Mnx1*) expression was largely restricted to SC and CN12 motor neurons. Glial markers, *Gfap*, *Mfge8*, *Sox10*, *Pdgfrb*, *Enpp6* and *Mog* were detectable in our neuron samples, but at much lower levels than in glial populations, while the macrophage/microglia markers *Itgam (CD11b)* and *Ccl3* were absent from our neuronal samples (Supplemental Fig. S2C). This cross-comparison demonstrated that our LCM-seq samples were highly enriched in neuronal transcripts and only included minor contaminations of glial transcripts. We analyzed the sustained homeobox transcription factor (Hox) gene profiles of collected neurons (Hedlund et al. 2010; Nichterwitz et al. 2016), which confirmed their positions along the anterior-posterior body axis. RN and CN3/4 neurons, which are located in the midbrain were, as expected, devoid of Hox gene expression (Supplemental Fig. S2D). Together, these data confirmed the identity, purity and high quality of our neuronal samples.

To investigate reproducibility among biological replicate samples and to determine correlations between different neuron types we conducted pairwise Spearman correlation. This analysis demonstrated a high correlation between samples originating from the same neuron type and revealed a close relationship between SC, CN7 and CN12 motor neurons (Fig. 1B). After unsupervised hierarchical clustering, midbrain neurons (RN and CN3/4) were distinct from other brainstem and SC motor neurons, while visceral CN10 motor neurons clustered separately within this group. Principal component analysis (PCA) on the entire gene expression data set revealed clustering of cell types with a strong influence of their developmental origin (Fig. 1C). Consequently, CN3/4 motor neurons clustered closely to RN neurons consistent with their specification in the ventral midbrain (Deng et al. 2011). CN7 and CN12 motor neurons formed a dense cluster with SC motor neurons, while visceral CN10 motor neurons clustered distinct from all the other cell populations. Altogether, we could demonstrate that biological replicates show high reproducibility, while neuronal groups form distinct clusters indicative of a high sensitivity.

In-depth gene expression analysis across neuron types confirmed known marker gene expression and identified several novel, cell-type specific transcripts. We confirmed their mRNA localization in the adult central nervous system using the Allen Mouse Brain Atlas (© 2004 Allen Institute for Brain Science. Allen Mouse Atlas available at www.brain-map.org). We could show that *Cxcl13* was restricted to RN neurons (Supplemental Fig. S3A). The transcription factor *Lhx4* was preferentially expressed in CN3/4 motor neurons (Supplemental Fig. S3B), while *Shox2* was present in CN7 motor neurons (Supplemental Fig. S3C). The peptide hormone *Nppb* distinguished visceral CN10 motor neurons from the other cell types (Supplemental Fig. S3D), while the proteoglycan *Dcn* was a marker for CN12 motor neurons (Supplemental Fig. S3E). We could thus validate the cell-type specific expression from our RNA-seq analysis with available *in situ* data.

Spinal motor neurons display inefficient *SMN2* exon 7 splicing compared to other cells in the spinal cord, thus rendering these cells low in full-length SMN (Ruggiu et al. 2012). While the LCM-seq method is highly sensitive for expression analysis it shows a clear 3‘ bias in gene body coverage, thus not allowing for in-depth splicing analysis (Nichterwitz et al., 2016). Therefore, to investigate if *SMN2* splicing differences could account for the differential susceptibility among the neuron types investigated here, we examined the expression levels of full-length *SMN2* mRNA by qPCR. In the majority of samples, we could not detect any exon 7 inclusion and where we did, the signal was close to the detection threshold (Supplementary Fig. S2E). Thus, the differential vulnerability of the neuron types investigated is not explained by differences in exon 7 splicing efficiency. Our RNA sequencing data therefore warrants further investigation to identify cell intrinsic mechanisms of selective neuronal vulnerability in SMA.

### SMA mice do not present a general developmental delay as determined by gene expression and neuromuscular junction analyses

It has been described that SMA patients and transgenic mouse models display a developmental delay in their neuromuscular systems (reviewed in Hausmanowa-Petrusewicz and Vrbová 2005). Differences in the developmental state of SMA and control motor neurons that may not reflect a pathological mechanism *per se* could hamper the identification of disease-relevant transcriptional changes. We thus addressed this issue here using our gene expression data. We first used weighted gene correlation network analysis (WGCNA) to identify gene sets that were regulated during early postnatal development of control somatic motor neurons. We chose the three modules that were most highly correlated with the early (P2) and late (P10) time points and changed over time (from a negative to a positive correlation or vice versa) (Supplemental Fig. S4A, B, Supplemental Table S2). PCA with the total of 5,843 genes in these modules confirmed the separation of somatic motor neurons based on age along PC1 (Supplemental Fig. S4C) supporting a change in expression of these genes during normal postnatal development. Gene ontology (GO) analysis revealed enrichment of several terms related to development (Supplemental Fig. S4D), and we used the 1,341 genes belonging to these terms (Supplemental Table S2) to perform PCA of control and SMA samples. As expected, the PCA demonstrated a clear age component between P2 and P10 samples. However, SMA samples did not appear to cluster differently on the age axis (PC1) compared to control samples (Fig. 2A). To better visualize the position of the SMA samples along the “age component” we plotted each cell type separately along the first principal component (PC1). We did not observe any difference in the age component in P2 and P5 samples, while P10 SMA samples shifted towards the younger age (PC1 negative) relative to control samples (Fig. 2B). As this shift occurred only at a time when SMA mice are visibly affected by disease, it likely reflects a disease process rather than a general developmental delay. In support of this, only 10.1% (136 genes) of all developmentally regulated genes were also significantly differentially expressed in disease (Fig. 2C, Supplemental Table S2), confirming that the majority of age-regulated genes are not affected in SMA. Further, if there was indeed a developmental delay in SMA motor neurons, we would expect to find genes regulated in the opposite directions in development and disease. However, of the 71 genes with opposite regulation, 76% (54 genes) were only regulated at the latest time point in disease, supporting our findings from the PCA (Fig. 2A, B). Altogether, our data challenges the notion that SMA motor neuron somas display a general developmental delay.

**Figure 2.**
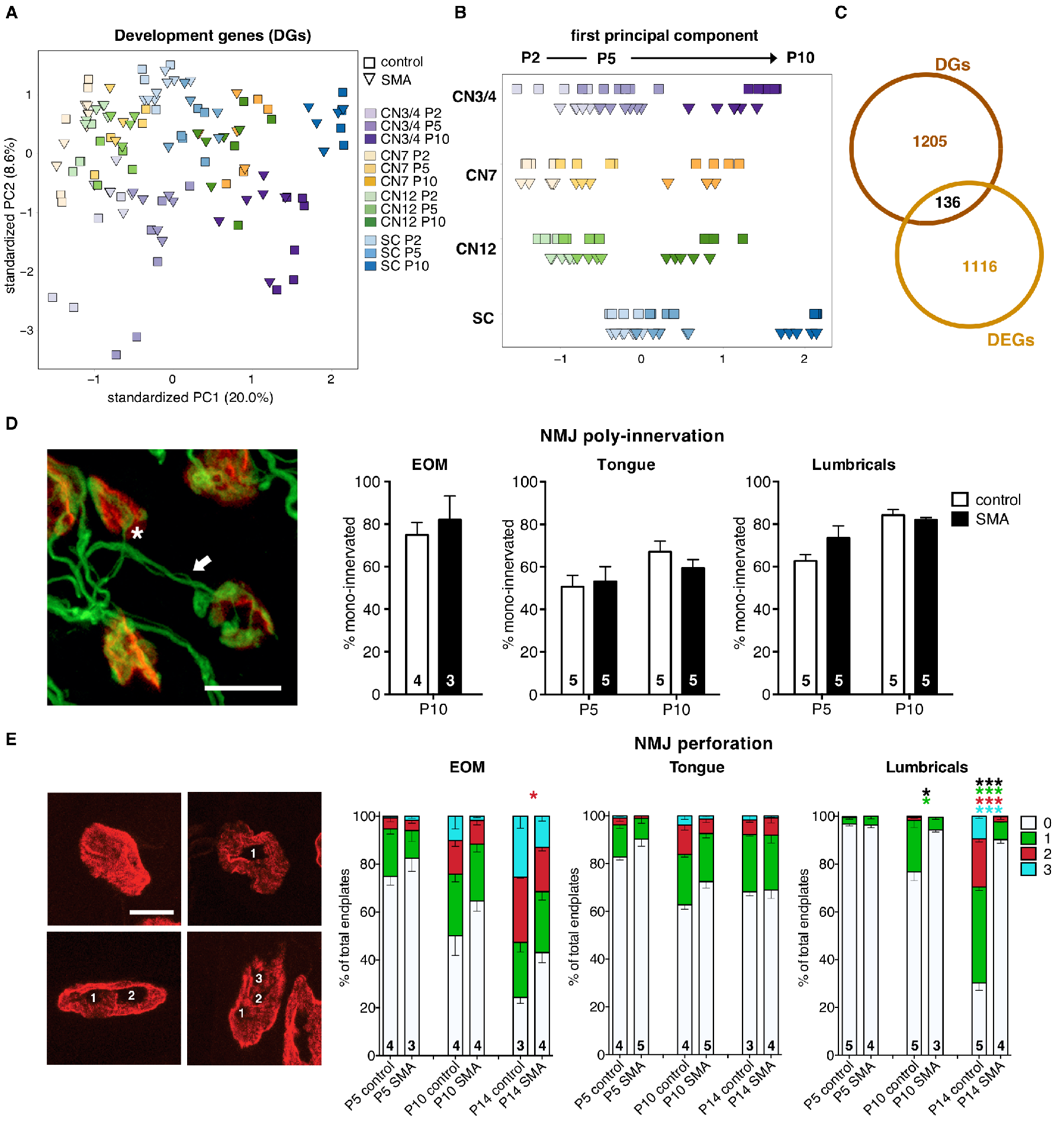
Evaluation of the developmental state of SMA somatic motor neuron somas and peripheral synapses. *(A)* PCA with development-related genes (DGs, 1,341 genes) of all somatic motor neurons in control and SMA. *(B)* Plot of PC1 alone to better visualize the “age component”. *(C)* Venn diagram depicting overlap between DGs and all differentially expressed genes between control and SMA (DEGs, no fold change cut-off, *P*_adj._<0.05) per cell type and time point. *(D)* Quantification of neuromuscular junction (NMJ) poly-innervation of extraocular muscles (EOM), tongue and lumbrical muscles. Micrograph shows examples of a mono-innervated (asterisk) and a poly-innervated (arrow) endplate. Red = alpha-bungarotoxin, post-synaptic nicotinic acetylcholine receptors; green = pre-synaptic, Neurofilament 165 kDa (2H3) and Synaptic vesicle protein 2 (SV2). *(E)* Quantification of NMJ perforation. Micrographs show examples of NMJs with 0, 1, 2 and 3 perforations. Red = alpha-bungarotoxin, post-synaptic nicotinic acetylcholine receptors. Numbers on bars in *D* and *E* represent the numbers of animals used for quantification. Multiple two-tailed t-tests per group (except for EOM in *D* where only one time point was investigated), * *P*_adj._<0.05, **** *P*_adj._<0.0001. Scale bar in *D* = 10 μm and in *E* = 10 μm (applicable to all micrographs in *E*).

To exclude any possible peripheral developmental phenotype in the neuromuscular system in SMA mice, we quantified the levels of poly-innervation and endplate perforations in target muscle groups. These measures are excellent indicators of early postnatal developmental stages until P14 in mice. Specifically, at neonatal stages muscle endplates are innervated by multiple incoming axons (polyinnervated). Through a process termed synaptic pruning the majority of these inputs are removed to leave NMJs mono-innervated by two weeks of age (in rodents). As muscle endplates mature they transition from a plaque-like morphology to a pretzel form. Increasing numbers of endplate perforations (holes in the staining) therefore indicate a more complex architecture of more mature endplates. Our analysis of extraocular muscles (EOM, innervated by CN3/4 motor neurons), tongue muscles (innervated by CN12 motor neurons) and lumbrical muscles from the hind limbs (innervated by lumbar SC motor neurons) demonstrated that the level of polyinnervation was equal in SMA mice and control littermates (Fig. 2D, multiple twotailed t-tests). We could also show that perforations were completely unaffected by disease in tongue muscle, and only very slightly affected in extraocular muscles at end-stage of disease (P14, multiple two-tailed t-tests, *P*adj.<0.05). As expected, lumbrical muscle endplates were severely affected at late stages of disease, lacking in perforations compared to control muscles (Fig. 2E; multiple two-tailed t-tests, *P*adj.(P10)<0.05, *P*adj.(P14)<0.0001). Collectively, the NMJ poly-innervation and endplate perforation data demonstrate that there was no obvious developmental delay in the maturation of neuromuscular synapses in agreement with the transcriptome data.

We therefore conclude that the motor systems develop normally in SMA mice, but that these are affected as disease progresses. Consequently, the same ages in control and SMA mice can be compared to distinguish disease-induced transcriptional changes without developmental processes obscuring the data sets.

### SMA-induced gene expression changes imply a common p53-mediated stress response in vulnerable and resistant somatic motor neurons

Towards our main goal of elucidating mechanisms of neuronal resilience and susceptibility, we investigated the transcriptional dynamics in resistant and vulnerable neurons in SMA. Using DESeq2, we performed pairwise differential expression analyses between control and SMA per cell type and time point. Notably, we found the strongest early transcriptional response in resistant CN3/4 and CN12 motor neurons with 134 and 211 differentially expressed genes (DEGs; no fold change cutoff, adj. *P*-value <0.05) at P2, followed by resistant RN and CN10 neurons with 57 and 55 regulated genes, respectively. At the late disease stage (P10), CN3/4, CN7 and SC motor neurons demonstrated a large number of disease-regulated genes, while RN and CN10 neurons showed minimal gene expression changes (Fig. 3A, Supplemental Table S3). Hierarchical clustering using DEGs of all neuronal populations separated SMA samples from controls at the earliest disease stage analyzed (P2) (Supplemental Fig. S5A). At later disease stages (P5-P10) somatic motor neuron groups clearly clustered together based on genotype, while RN and CN10 neurons did not separate with disease (Supplemental Fig. S5B, C). This demonstrates that resistant CN3/4 motor neurons display a response to disease that is distinct from other resilient neuron groups.

**Figure 3.**
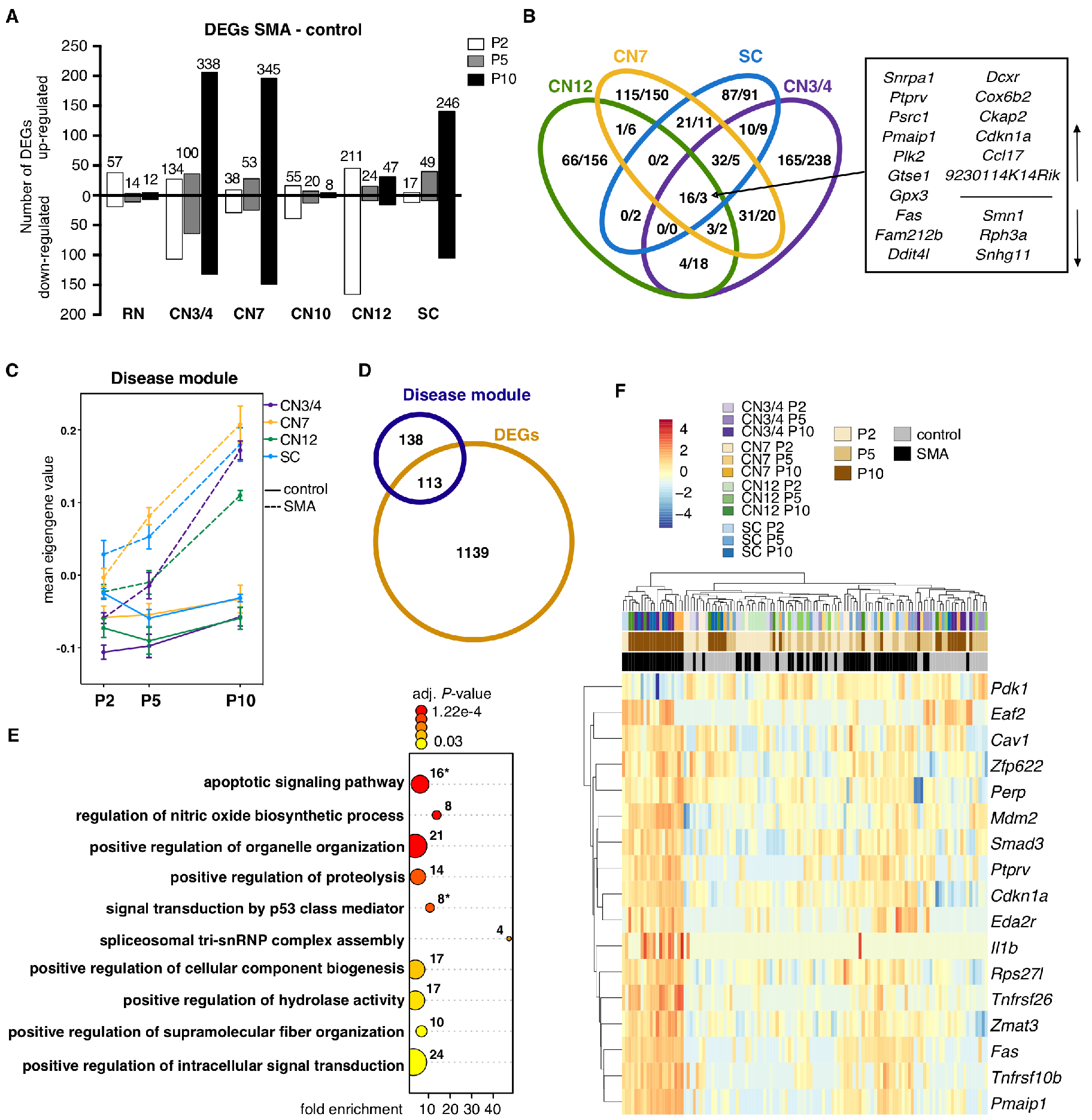
Analysis of disease-induced gene expression changes in SMA somatic motor neurons. *(A)* Number of significant genes from pairwise differential expression analysis per cell type and time point (DEGs, no fold-change cut-off, *P*_adj._<0.05). Numbers on bars represent total numbers of DEGs. *(B)* Venn diagram depicting the overlap in gene expression changes between somatic motor neurons (number of up-/down-regulated genes in SMA, all time points combined). *(C)* Mean eigengene values (first principal component of the disease module) within replicates. *(D)* Venn diagram depicting the overlap between genes in the disease module and DEGs. *(E)* GO term analysis for biological processes of the 251 genes in the disease module. Shown are selected GO terms, a complete list of enriched terms can be found in Supplemental Table S7. Numbers indicate the number of genes in a given term, color scale = adjusted *P*- value. Asterisks indicate gene sets that are plotted in *F. (F)* Expression heatmap of genes that belong to GO terms related to apoptosis and p53 signaling. Expression values were log_2_-transformed and mean centered.

To identify uniquely and/or commonly regulated genes across somatic motor neuron groups we plotted all DEGs in a Venn diagram. We only identified three commonly downregulated genes across all somatic motor neurons; *Smn1*, *Rph3a* and *Snhg11*. Sixteen transcripts were commonly upregulated with disease, including *Snrpa1, Plk2, Fam212b, Ddit4l, Dcxr, Cox6b2, Ccl17, 9230114K14Rik, Ptprv, Psrc1, Pmaip1, Gtse1, Gpx3, Fas, Ckap2 and Cdkn1a* (Fig. 3B). The majority of gene expression changes however, were unique to each neuron group (Fig. 3B, Supplemental Fig. S5D, Supplemental Tables S4 and S5) indicating distinct mechanisms to adapt to the loss of *Smn*.

To corroborate our DESeq2 analysis, we conducted WGCNA for all somatic motor neurons (both control and SMA samples), which revealed a module consisting of 251 genes that was highly positively correlated with disease and negatively correlated with control samples (Supplemental Fig. S5E, Supplemental Table S6), which we will refer to as “disease module”. As demonstrated by the mean eigengene values for the module within sample replicates, there was a clear genotype separation with disease progression in all somatic populations independent of their vulnerability (Fig. 3C). We plotted all genes of the module in a heatmap, which confirmed the similar expression levels across all somatic motor neurons (Supplemental Fig. S5F). Unaffected RN and CN10 neurons, on the other hand, did not show separation based on disease (Supplemental Fig. S5G). Besides *Smn*, only one gene of the disease module, *Plek2*, was differentially expressed in RN, while four genes, *Lars2*, *Olig2*, *Ptgds* and *Gm514*, were regulated in CN10 motor neurons with disease (Supplemental Fig. S5H). Furthermore, 45% of the genes (113 genes) in the module were identified as significantly differentially expressed with disease in one or more somatic populations using DESeq2 (Fig. 3D), including 14 of the 19 DEGs that are common to all somatic motor neurons. We thus identified a disease-signature specific to somatic motor neurons independent of their vulnerability. GO analysis for biological processes resulted in the significant (adj. *P*-value <0.05) enrichment of 23 GO terms in total (Fig. 3E, Supplemental Table S7). The pathways we identified as regulated included e.g. apoptotic signaling, signal transduction induced by p53 class mediator, positive regulation of intracellular signal transduction, spliceosomal tri-snRNP complex assembly, as well as several terms suggesting changes in metabolism (proteolysis, hydrolase activity) and cellular component organization (e.g. organelle organization, supramolecular fiber organization, cellular component biogenesis). To better visualize the timing of p53 and apoptosis marker expression in somatic motor neurons, we plotted the 17 dysregulated genes identified in these pathways in a heatmap (Fig. 3F). This analysis clearly demonstrated that activation of this pathway strengthened with disease progression and was shared by all somatic motor neuron populations investigated here. Consistently, GO term analysis of DEGs indicated a p53 pathway activation in SMA SC motor neurons already at P5 and confirmed the regulation of p53 and DNA damage pathways at P10 in SC, CN7 and CN3/4 motor neurons (Supplemental Table S7). Furthermore, eight of the 16 commonly upregulated genes from our DESeq2 analysis (Fig. 3B) are involved in the regulation of cell death, stress and/or p53 signaling. We used the STRING database to retrieve protein-protein interaction networks within the disease module. As expected, the network confirmed a coordinated p53-pathway activation. We further revealed a sub-network of genes involved in RNA processing and splicing, consistent with the role of SMN in spliceosome assembly (Supplemental Fig. S6).

In summary, we reveal a potentially detrimental common disease signature in all somatic motor neurons that is absent in RN and CN10 neurons. Notably, ocular motor neurons presented a unique adaptation mechanism to the loss of *Smn* that warranted further investigation.

### Resistant motor neurons activate a unique transcriptional program that could confer protection in SMA

To understand why CN3/4 motor neurons remain resilient to degeneration in SMA, while apoptotic signaling pathways appear activated, we conducted a close comparison with vulnerable SC motor neurons. Both neuron groups displayed distinct temporal responses (Fig. 4A, Supplemental Table S8). There was no overlap between CN3/4 and SC motor neurons in their early response to loss of *Smn* (Fig. 4B, Supplemental Table S9). Interestingly, 43% of all CN3/4- regulated genes at P2 belong to the GO term ‘nucleus’ including several genes involved in RNA processing and transcriptional activation/repression, such as *Rqcd1*, *Ube2b*, *Wtap*, *Cbx6*, *Hdac6*, *Ino80* and *Jmjd1c* (Supplemental Table S7) suggesting an early fine-tuning of transcription. As disease progressed, more genes were jointly regulated across the neuron groups, but the majority of transcriptional changes were still unique to each neuron type. Specifically, at P5, three genes were commonly upregulated in CN3/4 and SC motor neurons, while at P10, 56 genes were commonly upregulated and 11 downregulated (Fig. 4B). Thus, 100% of the genes regulated at P2 were unique to CN3/4 motor neurons, 97% at P5 and 80% at P10.

**Figure 4.**
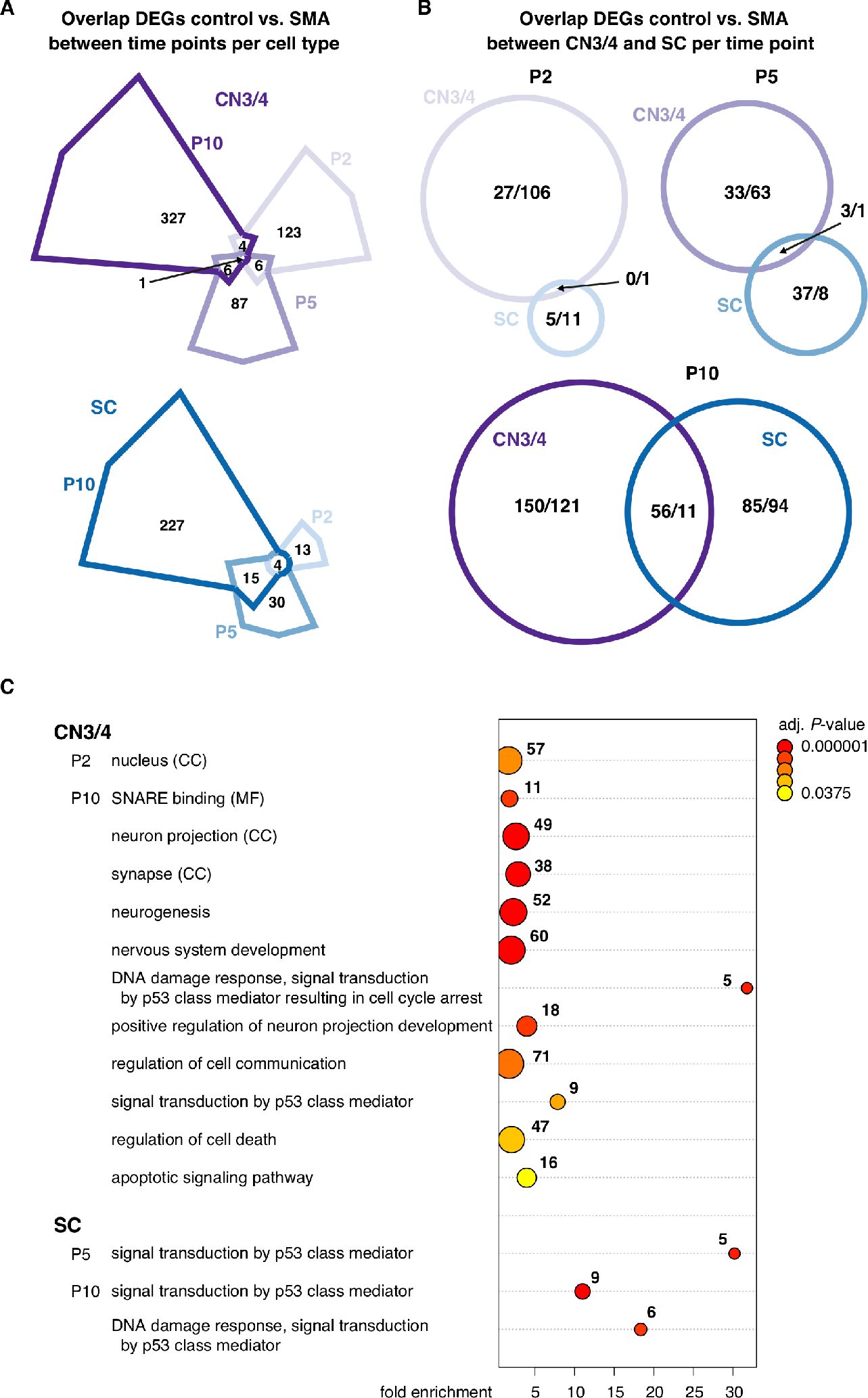
Comparison of gene expression changes in ocular and spinal motor neurons. *(A)* Venn diagrams depicting shared and time point specific DEGs between control and SMA motor neurons in CN3/4 (top) and SC (bottom). *(B)* Venn diagrams illustrating the overlap of DEGs between CN3/4 and SC at each time point. *(C)* GO term analysis of DEGs in CN3/4 and SC per time point. Shown are selected terms, a complete list of enriched terms can be found in Supplemental Table S7. Numbers indicate the number of genes in a given term, color scale = adjusted *P*- value. Terms belong to the domain biological processes unless specified otherwise; CC = cellular compartment, MF = molecular function.

GO term analysis of DEGs in CN3/4 neurons at P10 pinpointed a number of fundamental processes that were activated in response to disease. These pathways included neurogenesis, nervous system development, positive regulation of neuron projection development, regulation of cell communication, regulation of apoptotic processes and cell death (Fig. 4C, Supplemental Table S7). Among the enriched cellular compartments were neuron projection and synapse. To visualize CN3/4-restricted pathways that could be protective we used the STRING database to retrieve protein-protein interaction networks from all DEGs at P10. We obtained a highly interconnected protein network that consisted of 158 genes, corresponding to 46.7% of all DEGs in CN3/4 at P10 (Fig. 5) thus indicating a highly coordinated transcriptional response. The respective network for DEGs in SC motor neurons consisted of only 61 genes (25.5% of all DEGs), and a second major network included 22 genes (8.9% of all DEGs) (Fig. 6). P53 *(Trp53)* and 15 of the directly interacting proteins were upregulated in both neuron types (Fig. 5 and Fig. 6, grey outlines), in line with our GO term analysis. Upregulated genes involved in DNA damage repair, were *Polk*, (shared), *Tnks2* and *Mgmt* (CN3/4-specific) and *Rad51d* and *Timeless* (SC-specific). Importantly, CN3/4-specific downregulation included pro-apoptotic factors like *Itpr1* and *Dffa*, which was accompanied by the upregulation of anti-apoptotic and survival factors such as *Pak4*, *Pak6*, *Chl1*, *Tmbim4*, *Aldh4a1* and *Gdf15*. Neurotransmitter release is impaired in motor nerve terminals in SMA (Ruiz and Tabares, 2014). It is therefore compelling to see the CN3/4-specific regulation of genes involved in neurotransmitter release, including the upregulation of *Syt1*, *Syt5* and *Cplx2*, suggesting a compensatory mechanism in the disease-resistant cells. Among the many DEGs that are implicated in cytoskeletal reorganization, we found CN3/4-specific regulation of genes that are important for neurite outgrowth including *Gap43*, *Chl1*, *Syt1*, *Cald1* and *Serpine2*. (Fig. 5, 7). In contrast, in vulnerable SC motor neurons, we found increased levels of the actin filament severing *Inf2* (Supplemental Table S3) and the tubulin isoform *Tubb6*, which is associated with decreased microtubule stability (Bhattacharya and Cabral 2004; Salinas et al. 2014), and a significant decrease in mRNA levels of several motor proteins (*Dnahc2*, *Kif3a*, *Kif5a*) including genes that function in the anterograde transport of a variety of cargo to the cell periphery including the synapse (Fig. 6, 7).

**Figure 5.**
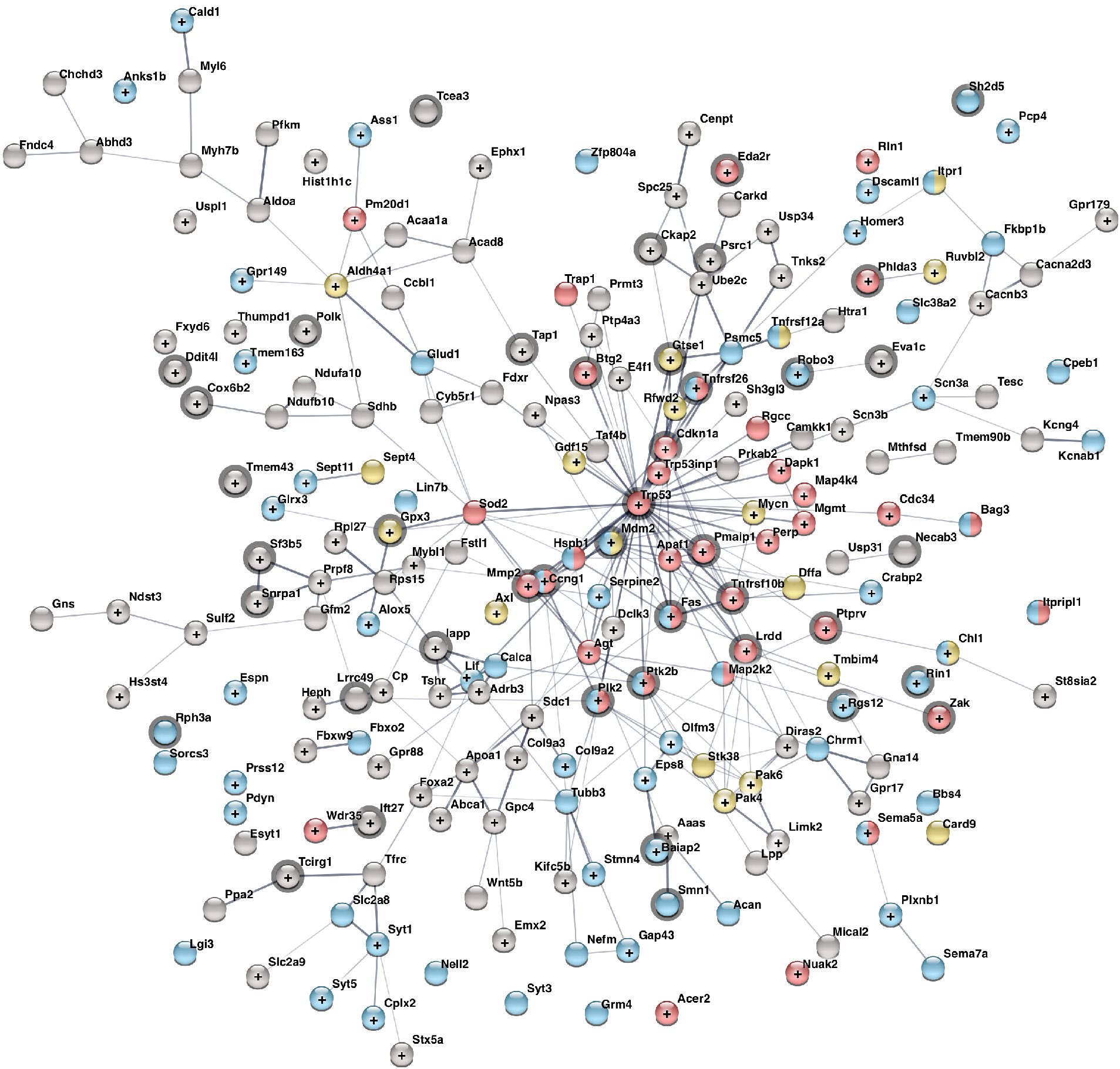
Protein-protein interaction network of DEGs in ocular motor neurons at P10. STRING analysis for protein-protein interactions including all 338 DEGs between control and SMA in ocular motor neurons at P10. Nodes are color-coded based on selected GO terms shown in Fig 4C: (red) signal transduction by p53 class mediator and regulation of cell death; (blue) positive regulation of neuron projection development, neuron projection and synapse; (yellow) downregulated apoptotic and upregulated survival genes. (+) upregulated genes in SMA, (empty nodes) downregulated in SMA. (Grey outlines) genes that are also differentially expressed in SC neurons. Network edges represent the confidence of the predicted associations between nodes (edge thickness = strength of data support). Disconnected nodes that do not belong to any of these GO terms are not shown.

**Figure 6.**
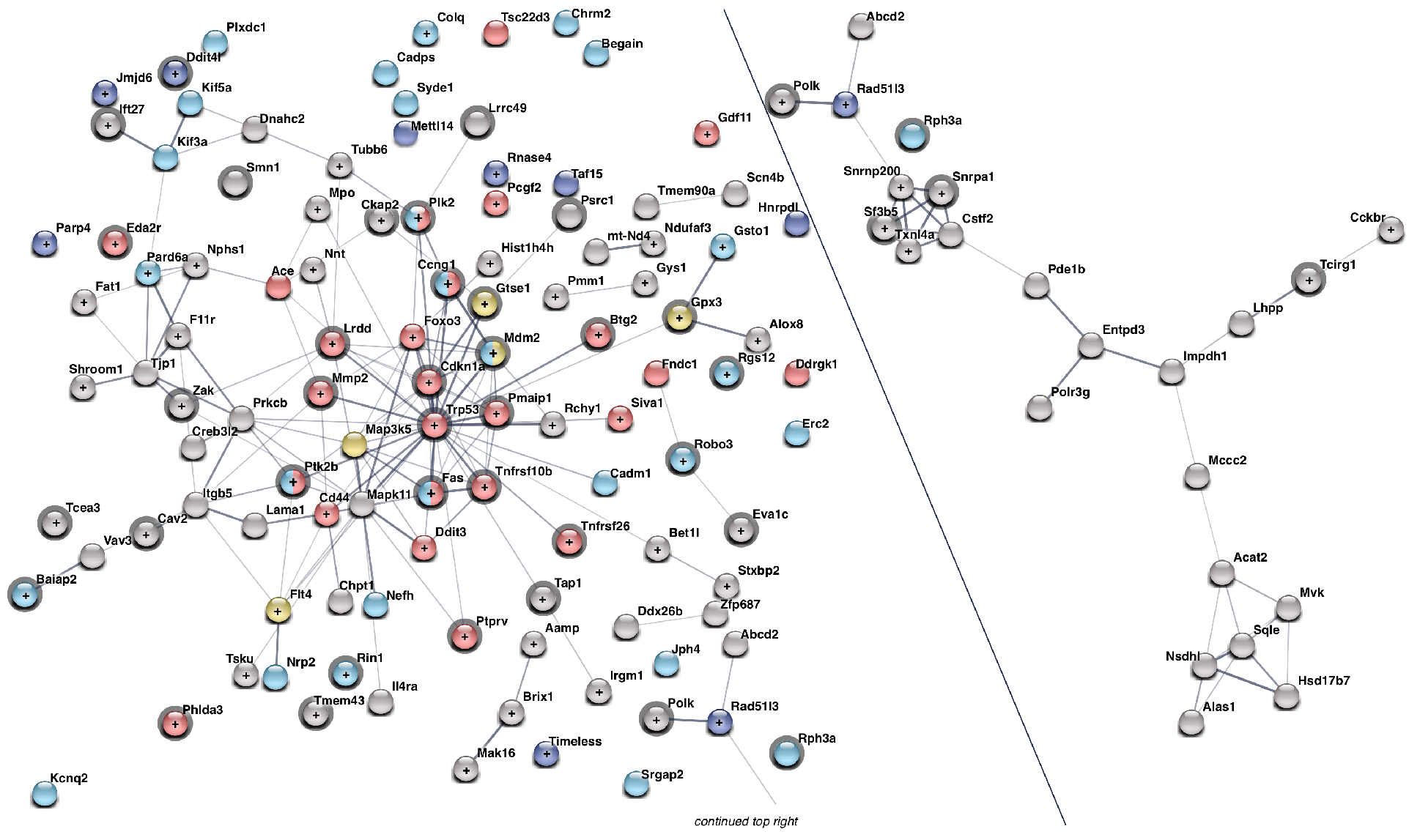
Protein-protein interaction networks of DEGs in spinal motor neurons at P10. STRING analysis for protein-protein interactions including all 246 DEGs between control and SMA in spinal motor neurons at P10. Nodes are color-coded based on selected GO terms shown in Fig 4C including CN3/4- specific terms: (red) signal transduction by p53 class mediator and regulation of cell death; (blue) positive regulation of neuron projection development, neuron projection and synapse; (yellow) downregulated apoptotic and upregulated survival genes. (+) upregulated genes in SMA, (empty nodes) downregulated in SMA. (Grey outlines) genes that are also differentially expressed in CN3/4 neurons. Network edges represent the confidence of the predicted associations between nodes (edge thickness = strength of data support). Disconnected nodes that do not belong to any of these GO terms are not shown except for selected genes (dark blue).

**Figure 7.**
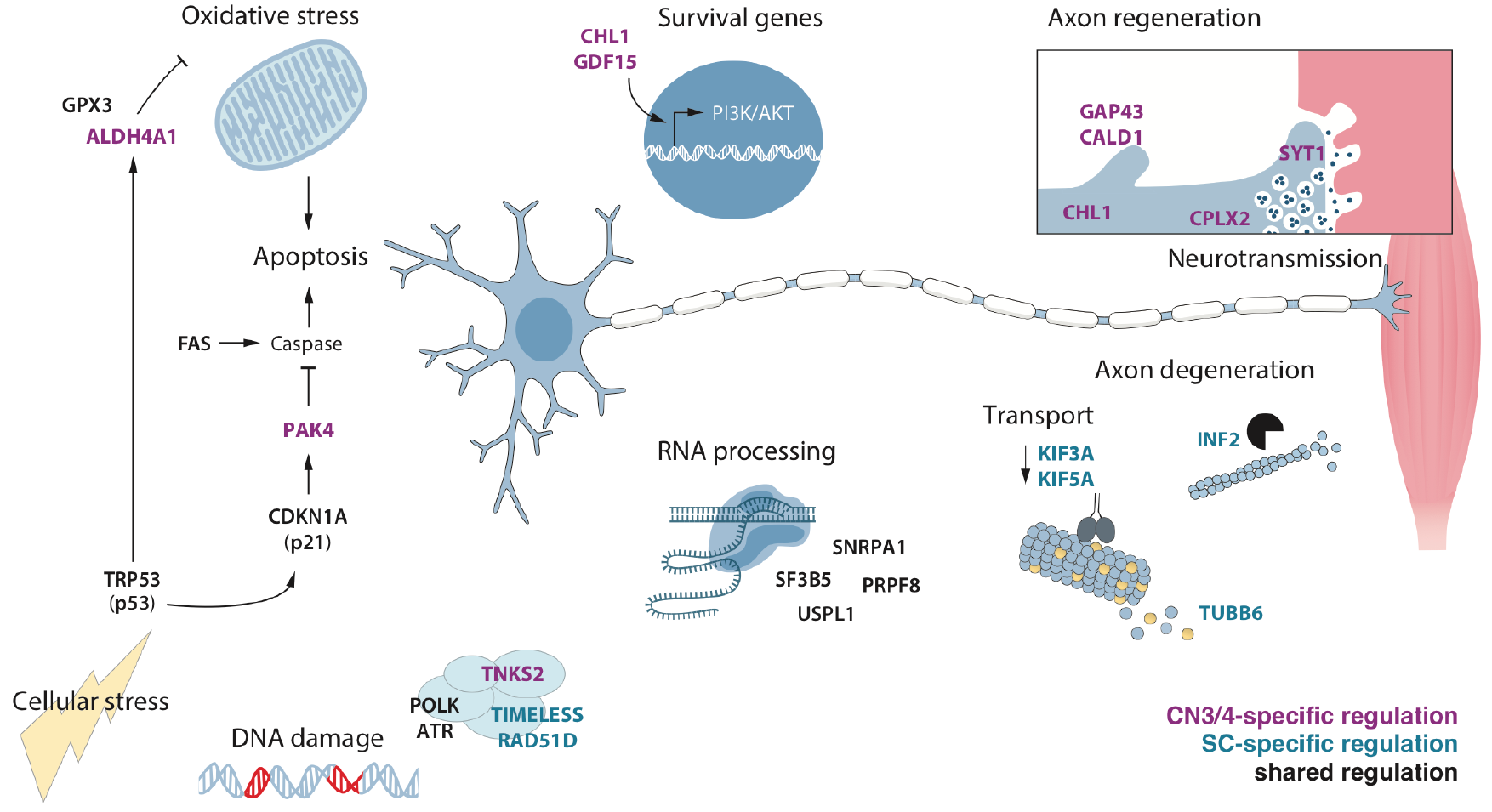
Common and cell-type specific disease mechanisms in SMA. Somatic motor neurons display transcriptional changes due to the loss of full length SMN protein that are distinct from red nucleus and vagus neurons. For example, prominent changes in expression levels of genes that function in RNA processing are restricted to somatic motor neurons. These neurons are furthermore exposed to cellular stress including oxidative stress and DNA damage, and DNA repair genes are induced. P53- and cell death signaling pathways are activated in all somatic motor neurons independent of their susceptibility to degeneration in SMA. Vulnerable spinal (SC) motor neurons show signs of axon degeneration and axonal transport deficits. Resistant ocular (CN3/4) motor neurons selectively upregulate the expression of genes that counteract apoptosis and promote cell survival. Increased levels of genes functioning in neurite outgrowth, axon regeneration and neurotransmission, which support the maintenance of a functional neuromuscular synapse, are also seen in ocular motor neurons in disease.

Collectively, our analysis shows that resistant neurons respond early and uniquely to the loss of *Smn*. Arguably this response involves the regulation of *bona fide* neuroprotective factors and processes.

## Discussion

In SMA, somatic motor neurons are selectively vulnerable to the loss of the broadly expressed *SMN1* gene while certain somatic motor neuron groups, including oculomotor and trochlear neurons, are for unknown reasons relatively resistant to degeneration. In this study, we conducted a longitudinal analysis of the transcriptional dynamics in resistant (CN3/4 (ocular), CN10 (vagus), CN12 (hypoglossal) and RN (red nucleus)) and vulnerable (CN7 (facial) and SC (spinal)) neuron groups during disease progression in SMA mice. We found that, independent of their vulnerability, somatic motor neurons activate p53 signaling pathways that are associated with DNA damage and cell death. This upregulation was absent in visceral vagus motor neurons and red nucleus neurons. This suggests that the activation of the p53 pathway is a stress response specific to somatic motor neurons in SMA but that it does not necessarily lead to degeneration. We also show that the majority of gene expression changes induced by the loss of *Smn* are cell-type specific and reveal several pathways that are restricted to resistant ocular motor neurons. Such *bona fide* protective pathways include increased levels of survival factors, while pro-apoptotic genes are selectively downregulated (Fig. 7). Importantly, we observed ocular-motor restricted transcriptional regulation of genes involved in neurotransmission and neurite outgrowth that may aid in the maintenance of a functional neuromuscular synapse and thus contribute to selective resistance in SMA motor neurons.

Our initial analysis of neuronal populations showed that ocular motor neurons had a transcriptional profile that was distinct from other somatic motor neuron groups. This transcriptional distinction of ocular motor neurons is consistent with previous microarray studies conducted on tissues from control rats (Hedlund et al. 2010), mice (Kaplan et al. 2014) and humans (Brockington et al. 2013). Our PCA also revealed that the global gene expression of ocular motor neurons was more similar to other somatic motor neuron groups and red nucleus neurons than to visceral vagus motor neurons. Red nucleus and ocular neurons display commonalities as these cell types share a developmental origin and are both specified from *Lmx1b*-positive floor plate progenitors, located lateral to the dopamine domain (Deng et al. 2011). Ocular motor neurons share a developmental expression of the transcription factors *Islet1/2*, *Phox2a* and *Phox2b* with other cranial motor neuron groups, including facial and hypoglossal. However, ocular motor neurons uniquely lack the expression of *Hb9* (*Mnx1*) and are not subjected to patterning from Hox genes during development, unlike all other motor neuron groups in the brainstem and spinal cord (reviewed in Cordes 2001). Hox genes are important for establishing motor neuron pool identity as well as connectivity with muscle targets (Dasen et al. 2005) and as such it is not surprising that ocular motor neurons are distinct from other somatic motor neuron groups. Nonetheless, based on the entire gene expression profile, ocular motor neurons were more closely related to somatic motor neurons than to visceral vagus motor neurons.

Additionally, we detected cell-type specific transcripts in several neuronal populations. *Shox2* is a transcription factor involved in pattern formation. We found *Shox2* expression restricted to facial motor neurons in line with the finding that *Shox2* is required for proper development of the facial motor nucleus (Rosin et al. 2015). While *Dcn*, coding for the proteoglycan decorin, was to some degree detectable in facial motor neurons, *Dcn* levels in hypoglossal motor neurons were strikingly higher at all time points investigated, revealing this gene as an excellent marker for the hypoglossal nucleus. To the best of our knowledge, there was previously no established marker available that reliably distinguishes hypoglossal from other brainstem motor neurons. Finally, we found *Cxcl13* to be highly restricted to red nucleus neurons, which adds to the existing repertoire of markers for this midbrain nucleus. Thus, we have revealed cell-type specific markers that can be highly valuable tools to selectively modulate cellular functions and identify distinct cell types *in vitro* and *in vivo*.

The simplest explanation for differential neuronal vulnerability in SMA would be the preferential inclusion of exon 7 in SMN2 transcripts in resilient neuronal populations and thus presence of higher levels of full-length SMN protein. However, using qPCR, we found that the splicing of the *SMN2* gene itself was consistent across neuron types, with minimal inclusion of exon 7 in both resistant and vulnerable neurons. Thus, differential splicing of *SMN2* is not the cause of differential neuronal vulnerability in this model.

Our analysis of motor neurons in SMA revealed a striking upregulation of p53 signaling across all somatic motor neuron groups in disease. P53 is a master regulator in response to several cellular stressors such as oxidative stress or DNA damage. SMN directly interacts with p53 (Young et al. 2002) and transcriptional activation of p53 or its target genes has been previously demonstrated in different models of SMA (Jangi et al. 2017; Murray et al. 2015; Staropoli et al. 2015; Zhang et al. 2013, 2008; Bäumer et al. 2009; Simon et al. 2017). Surprisingly, cell death signaling was not restricted to vulnerable populations in our study, but was also evident in resistant neurons, predominantly at a late stage of disease (P10). Consistently, more resistant spinal motor neurons of the lateral motor column show increased p53 protein levels with progression of disease (Simon et al. 2017). Importantly, p53 itself can activate a number of targets that exert anti-apoptotic effects, such as *Gtse1*, *Dcxr*, *Gpx* and *Cdkn1a* (reviewed in Jänicke et al. 2008), which were also upregulated in all our somatic populations. Specifically, the Cyclin Dependent Kinase Inhibitor 1A (*Cdkn1a*, also known as p21) plays a crucial role in cell cycle regulation and response to DNA damage. *Cdkn1a* can also prevent apoptosis, for instance by transcriptional repression of pro-apoptotic genes or inhibiting caspases (reviewed in Gartel and Tyner 2002). Thus, upregulation of *Cdkn1a* could be part of a protective response in somatic motor neurons, which is not sufficient to protect the most vulnerable cells.

Consistent with the activation of p53-signaling, our analysis suggests that DNA damage pathways are activated in this disease. This is also in line with previous reports investigating SMA-induced transcriptional changes (Jangi et al. 2017; Murray et al. 2015; Staropoli et al. 2015; Bäumer et al. 2009). However, direct evidence for DNA damage in SMA motor neurons is as yet limited. Three recent studies failed to detect increased *γ*H2AX (H2AX phosphorylated on serine 139) immunoreactivity, a reliable marker for DNA double strand breaks, in motor neurons in SMA mouse tissues and human SMA cells (Simon et al. 2017; Jangi et al. 2017; Murray et al. 2015). Interestingly, Tisdale et al. (2013) observed reduced levels of H2AX mRNA after Smn-knockdown and correspondingly lower induction of H2AX phosphorylation upon DNA damage, presenting a possible explanation for the lack of increased γH2AX in SMA motor neurons in the presence of DNA damage. Furthermore, knockdown of *Fus*, which can directly interact with SMN (Yamazaki et al. 2012), led to diminished γH2AX immunoreactivity in response to a genotoxic reagent, while DNA damage was validated using the comet assay (Wang et al. 2013). Our data support the presence of DNA damage as evidenced by increased expression levels of multiple genes with roles in DNA repair: *Atr*, which is a central player in the DNA damage response, is a member of the disease module that we found to be commonly regulated in all somatic motor neurons; the DNA polymerase *Polk* was upregulated in ocular, facial and spinal motor neurons; selectively upregulated transcripts in ocular motor neurons were the DNA repair gene O-6 methylguanine-DNA methyltransferase *Mgmt*, and the tankyrase *Tnks2*, which plays a role in RAD51 recruitment to DNA double strand breaks (Nagy et al. 2016); the RAD51 family member *Rad51d* was upregulated in spinal motor neurons as was *Timeless*, which can mediate DNA damage repair of double strand breaks (Young et al. 2015; Xie et al. 2015). It has been suggested that SMN is important for preventing DNA damage as it associates with RNA pol II (Pellizzoni et al. 2001) and is required for resolving RNA-DNA hybrids during transcription termination (Zhao et al. 2015). These findings provide a putative mechanism by which DNA damage could occur in SMN deficient cells. Furthermore, several genes involved in RNA regulation, including *FUS* and TDP-43, which when mutated cause the motor neuron disease amyotrophic lateral sclerosis (ALS), contribute to the prevention or repair of transcription-associated DNA damage. Depletion of FUS or TDP-43 leads to an increase in DNA damage (Hill et al. 2016) and it appears that loss of SMN could have a similar result. Notably, while p53 signaling and especially *Cdkn1a* levels were increased during pre-and early symptomatic disease stages (P2 and P5), expression levels of DNA repair genes were predominantly altered later in disease (P10). Thus, further investigation is warranted to elucidate if DNA damage is a major driver of neuromuscular pathology in SMA.

With our comprehensive comparison of resistant ocular and vulnerable spinal motor neurons using gene ontology and protein-network analyses, we were able to pinpoint several protective pathways that are selectively regulated in our resilient motor neuron group and thus likely counteract commonly activated stress responses. Prosurvival factors with increased levels in ocular motor neurons were for instance *(i)* the CDKN1A activated kinases *Pak6 and Pak4. Pak4* prevents caspase activation and thus apoptosis (Gnesutta and Minden 2003; Gnesutta et al. 2001) and encouragingly, its neuroprotective function was recently demonstrated in a rat model of Parkinson’s disease (Won et al. 2016). *(ii)* The mitochondrial aldehyde dehydrogenase *Aldh4a1*, which can safeguard cells from oxidative stress (Yoon et al. 2004). There is recent evidence for increased oxidative stress linked to mitochondrial dysfunction in primary motor neuron cultures of SMA mice (Miller et al. 2016). The enrichment of genes related to an oxidative stress response in the relatively vulnerable facial motor neurons (Supplemental Table S7) supports a role for oxidative stress in the pathology of SMA. *(iii)* The Growth differentiation factor 15 *(Gdf15)* is important for motor neuron survival (Strelau et al. 2009). *(iv) Tmbim4*, also known as Golgi anti-apoptotic protein (GAAP) or Lifeguard4, which exhibits anti-apoptotic functions likely through modulation of intracellular Ca^2+^ (reviewed in Carrara et al. 2017). Consistently, the Ca^2+^ channel *Itpr1* (IP3 Receptor), which plays a role in endoplasmic reticulum (ER) stress-induced apoptosis, was selectively downregulated in ocular motor neurons. The activation of ER stress pathways in SMA was demonstrated in a transcriptomics study using iPSC-derived motor neurons from SMA patients (Ng et al. 2015). Thus, resistant ocular motor neurons appear to, by necessity, regulate pathways that counteract the detrimental processes that are activated with disease in all somatic motor neurons. Other resistant neurons groups, including red nucleus and visceral motor neurons showed no such gene regulation, but also lacked DNA damage and p53 activation.

As SMN also functions locally in the axon, including nerve terminals, preventing apoptosis is unlikely to fully rescue motor neuron function. We therefore examined transcriptional regulation of genes involved in neurotransmission and neurite outgrowth, that would affect neighboring neurons and muscle in addition to motor neurons themselves. Exciting candidates that were specifically upregulated in ocular motor neurons were synaptotagmin 1 and 5 (*Syt1* and *Syt5*) (Fig. 7). SYT1 functions in the release of synaptic vesicles and has recently been associated with differential vulnerability in SMA (Tejero et al. 2016). By counteracting the impaired neurotransmitter release that has been observed in SMA motor neurons (Ruiz and Tabares 2014; Kariya et al. 2008), ocular motor neurons may be able to maintain their connection to target muscles and ensure their functionality. In support of this, we found complexin II (*Cplx2*) upregulated in ocular motor neurons, which also modulates synaptic vesicle release (Ono et al. 1998). The genetic depletion of *Cplx2* in mice results in locomotor deficits (Glynn et al. 2003) suggesting an important function in motor neurons. We also identified caldesmon 1 *(Cald1)*, the growth associated protein 43 (*Gap43*) and the L1 cell adhesion molecule homolog *Chl1* to be selectively upregulated in ocular motor neurons. CALD1 is a regulator of neurite outgrowth (Morita et al. 2012) and GAP43 is crucial for developmental axon outgrowth as well as regeneration. Likewise, CHL1 levels increase in regenerating motor neurons after sciatic nerve injury (Zhang et al. 2000) and it is a survival factor for primary rat motor neurons, acting via the PI3K/Akt pathway (Nishimune et al. 2005). It was recently shown that *Gap43* mRNA and protein levels were reduced in axons and growth cones of primary spinal motor neurons isolated from a severe mouse model of SMA (Fallini et al. 2016), predisposing these to a lower degree of re-connectivity. Thus, the SMA-induced increase in *Cald1*, *Gap43* and *Chl1* in ocular motor neurons could help these to reconnect to muscle targets if disconnected during disease.

In summary, the transcriptional regulation of genes related to neurotransmission and neurite outgrowth presents a compelling *bona fide* protective mechanism that is activated during disease progression selectively in these resistant motor neurons.

In conclusion, our study provides important insights into mechanisms of selective resistance and vulnerability in SMA. We show that all somatic motor neurons, irrespective of their vulnerability in SMA, present stress responses due to SMN deficiency. However, resistant ocular motor neurons selectively activate survival pathways and show transcriptional regulation of genes that are important for the maintenance and/or regeneration of a functional neuromuscular synapse. The modulation of such mechanisms presents a promising strategy not only for the additional treatment of SMA patients where splicing correction of *SMN2* is not sufficient but also other motor neuron diseases like ALS. We thus revealed novel targets that will be exciting to investigate further in the context of motor neuron disease.

## Methods

### Ethics statement and animal model

All animal procedures were approved by the Swedish ethics council and carried out according to the Code of Ethics of the World Medical Association (Declaration of Helsinki). Animals were housed under standard conditions with a 12 hour dark/light cycle and had access to food and water *ad libitum*. Neonatal pups were used as a model of SMA (*Smn*^−/−^/*SMN2*^+/+^/*SMNΔ7*^+/+^) and age matched littermates that were homozygously wild type for murine *Smn* (*Smn*^+/+^/*SMN2*^+/+^/*SMNΔ7*^+/+^) were used as controls (Le et al. 2005) (Jackson Laboratory stock number 005025). For transcriptomics, we used 2-, 5- and 10-day old pups (P2, P5 and P10), whereas neuromuscular junction analysis was performed on 5-, 10- and 14-day old pups (P5, P10 and P14). P2 and P5 pups were sacrificed by decapitation, while P10 and P14 pups were anesthetized with a lethal dose of avertin (2,2,2- Tribromoethanol in 2-Methylbutanol, Sigma-Aldrich) prior to decapitation.

### Reagents

For all RNA-seq experiments, nuclease-free water (H_2_O, LifeTechnologies) was used and all reagents were of molecular biology/PCR grade if available. Only tubes that were certified nuclease-free were used. All workbenches and equipment were cleaned with RNaseZAP (Ambion) and additionally with DNAoff (Takara) for library preparations. For the preparation of the different staining components prior to laser capture microdissection (LCM), 99.7% EtOH Aa (Solveco) and nuclease-free H_2_O were used.

### Laser capture microdissection of distinct neuronal populations

Brain and lumbar spinal cords were dissected and immediately snap frozen in 2-Methylbutane (Sigma-Aldrich) on dry ice. Tissue was stored at −80°C until further processing. Brains and in OCT embedded spinal cords were sectioned in a cryostat at −20°C (12 μm coronal sections) and placed onto PEN membrane glass slides (Zeiss) that were subsequently stored at −80°C until further processing. Motor neurons are easily identifiable by their large soma size and by their distinct locations in the ventral horn of the spinal cord and in the brainstem. Tissue was therefore subjected to a quick histological staining (Histogene, Arcturus/LifeTechnologies). Immediately prior to staining, slides were thawed for 30 seconds and subsequently placed into 75% EtOH for 30 seconds. After incubation for another 30 seconds in H_2_O, slides were incubated with 150-200 μl of Histogene staining solution for 20 seconds. Staining solution was removed by tapping the slide on lint-free tissue, followed by incubation for 30 seconds in H_2_O. Dehydration was achieved by incubating the slides in EtOH solutions of rising concentration (75%, 95% and 99.7% EtOH, 30 seconds each). Slides were then placed into the slide holder of the microscope and cells were captured using the Leica LMD7000 system. LCM was performed at 40x magnification and cutting outlines were drawn in close proximity to individual cells to minimize contamination by surrounding tissue. Only cells with an area of more than 200 μm^2^ (150 μm^2^ for vagus motor neurons) and a visible nucleus with nucleolus were selected. 100-200 cells were collected into the dry cap of 0.2 ml PCR soft tubes (Biozym Scientific). 5 μl lysis buffer (0.2% Triton X-100, with 2 U/μl recombinant RNase inhibitor, Clontech) were added to the cap and mixed by pipetting up and down 5 to 10 times. Samples were spun down using a table centrifuge (VWR) for 10 seconds and snap frozen on dry ice. The duration from thawing the slides until freezing of the sample never exceeded 2 hours. Samples were stored at −80°C until further processing.

### cDNA and sequencing library preparation

Library preparation was performed with a modified version of the Smart-seq2 protocol (Picelli et al. 2014b, 2013) and is described in detail in (Nichterwitz et al. 2018, 2016). For reverse transcription (RT), polymerase chain reaction (PCR) cycles and heat incubation steps a T100 Thermal Cycler was used (BioRad). RT was followed by 18 cycles of PCR amplification. After purification with magnetic beads (GE Healthcare), cDNA concentration and quality of cDNA libraries were determined with an Agilent 2100Bioanalyzer using the High Sensitivity DNA kit. For the generation of sequencing libraries, 1 ng of cDNA was used (as determined with Bioanalyzer, 100-9000 bp range). The tagmentation reaction was carried out with 0.4-1 μl of in house Tn5 (Picelli et al. 2014a). Ligation of sequencing indices (Nextera XT Sequencing Index Kit, Illumina) and enrichment PCR (10 cycles) was performed with Kapa HiFi polymerase. Before pooling the sequencing libraries, a purification step with magnetic beads was performed and the concentration of each sample was determined on a Qubit fluorometer (ThermoFisher) with the dsDNA high sensitivity kit (LifeTechnologies). An equal amount of cDNA from 15 to 30 samples was pooled per lane of a flow cell. Single read 43-bp sequencing was performed on an Illumina HiSeq 2000 sequencing system.

### RNA-seq data analysis

The RNA-seq reads were mapped simultaneously to the mm10 mouse genome assembly and the genomic sequence of human *SMN2* (including introns) from the hg19 assembly using STAR (version 2.4.1c) (Dobin et al. 2013). Quality control was performed with rrnaseq (available on GitHub at https://github.com/edsgard) to ensure sufficient sequencing depth and mapping ratios appropriate for our poly(A)-enriched sequencing strategy. We used uniquely mapped reads (69.1 ± 0.39% mean ± SEM, Supplemental Fig. S2B) for further analyses. Expression levels were determined using the rpkmforgenes.py software (available at http://sandberg.cmb.ki.se/rnaseq) with the Ensembl gene annotation. Only samples with more than 11,000 detected genes (≥1 RPKM) were included in the analysis. For principal component analysis and gene expression heatmaps, data was log_2_- transformed. Weighted gene correlation network analysis (WGCNA) (Langfelder and Horvath 2008) was performed using control somatic motor neuron samples to investigate developmental gene expression changes, and control and SMA somatic motor neurons for the analysis of SMA induced transcriptional changes. To eliminate expression noise, the gene sets included in the analyses were limited to genes expressed at RPKM ≥ 1 in at least three replicates of any sample type (i.e. consistently expressed in at least one group of cells with common genotype, cell type and time point). Module-trait correlations were calculated for categorical, binary traits. Examples of such “traits” could be cell type, genotype, timepoint or a combination of two or more of those categories. Each sample was given a value of 0 or 1 for each trait, depending on which group(s) it belongs to. Figure 3C and Supplemental Fig. S4B show the mean eigengene values (first principal component of the modules) within replicates. Differential expression was calculated using DESeq2 (Love et al. 2014) after first removing all genes with zero expression across all samples and adding a pseudocount of 1. The pseudo-count was added, as DESeq2 appeared to perform poorly when most replicates had a zero count. Independent filtering (independentFiltering = FALSE) and outlier filtering (cooksCutoff = FALSE) were disabled as their inclusion occasionally resulted in our control gene (*Smn*) lacking significance. Events were considered significant when the adjusted P-value was below 0.05. As the biological importance of a given change in expression level is unknown, no fold change cut-off was applied. Gene overlaps were calculated in R using the VennDiagram package (version 1.6.17). GO term analysis for biological processes, molecular function and cellular compartment was conducted with Panther (http://www.pantherdb.org/) (Mi et al. 2013) (Panther Overrepresentation Test, release 2017-04-13; GO Ontology database release 2017-09-26; Bonferroni correction of *P*-values). To retrieve protein-protein interaction networks, we used the STRING database (https://string-db.org, version 10.5 (Szklarczyk et al. 2017)) and uploaded selected gene sets using the ‘Multiple proteins’ query. The minimum required interaction score was set to 0.04 (medium confidence) and the interaction sources neighborhood and gene fusion were deselected. For the identification of genes that belong to certain GO terms (Fig. 6 and Supplemental Fig. S6) complete gene lists of these terms (filtered for *Mus musculus*) were downloaded from AmiGO 2 (http://amigo.geneontology.org/amigo) (The Gene Ontology Consortium 2017; Ashburner et al. 2000).

### Use of published datasets

For the evaluation of the purity of our samples, we compared our samples to a previously published dataset (Zhang et al. 2014). Raw data was obtained from the Gene Expression Omnibus (GEO, accession number GSE52564) and processed as described for our own samples.

### *SMN2* splicing analysis by quantitative PCR

For real-time quantitative PCR, 0.5 ng of Smart-seq2 libraries were used per 15 μL reaction with 500 nM of each assay primer (SMN2 total, 5’-GTG AGG CGT ATG TGG CAA AAT-3’, 5’-CAT ATA GAA GAT AGA AAA AAA CAG TAC AAT GAC C-3’; and SMN2 full-length, 5’-CAC CAC CTC CCA TAT GTC CAG ATT-3’, 5’-GAA TGT GAG CAC CTT CCT TCT TT-3’; Ruggiu et al. 2012) in 1x MESA Green qPCR Master Mix Plus (Nizzardo et al. 2014). The reactions and recording were performed on an Applied Biosystem 7500 Fast Real-Time PCR System in technical duplicates. The specificity of signals was controlled by recording dissociation curves following the amplification. CT values were extracted with the accompanying software and signals below lower detection limits and with unspecific amplification were deemed non-detected. The relative expression levels were calculated according to the ΔCT method (Pfaffl 2001).

### Neuromuscular junction analysis

For neuromuscular junction (NMJ) immunohistochemistry of extraocular, lumbrical (from the hind-paw) and tongue muscles, tissue was dissected in 0.1 M phosphate buffered saline (PBS, LifeTechnologies) and post-fixed for 30 and 60 min, respectively, in 4% paraformaldehyde (PFA, Sigma-Aldrich). Lumbrical muscles were left intact for whole-mount processing. To increase visible muscle surface, muscles were gently stretched laterally. Tongues were sectioned at 30 μm thickness. For visualization of NMJs, tissue was permeabilized in 4% Triton X-100 in 0.1 M PBS for 1 hour and blocked in 0.1% Triton X-100 in 10% donkey serum in 0.1 M PBS for 1 hour at room temperature. To label the NMJ and incoming axons, tissue was incubated with an anti-neurofilament (165 kDa) antibody (2H3, 1:50 (Dodd et al. 1988)) and an anti-Synaptic vesicle protein 2 antibody (SV2, 1:100 (Buckley and Kelly 1985)) at 4°C overnight. After washing in blocking solution for 1 hour at room temperature, muscles were incubated for 3 hours in Alexa Fluor 488-conjugated secondary antibody (1:500, LifeTechnologies), followed by washing in 0.1 M PBS for 30 minutes. Subsequently, muscle tissues were incubated with α-bungarotoxin (α-BTX) conjugated to tetramethyl-rhodamine isocyanate (TRITC, 1:1000; LifeTechnologies) for 10 min in order to visualize post-synaptic acetylcholine receptors (Comley et al. 2016). Muscles were whole-mounted on glass slides in Mowiol 4-88 (Sigma-Aldrich) and cover-slipped. Muscle preparations were imaged using a laser scanning confocal microscope (Zeiss LSM700 and LSM800). All images were acquired as Z-stacks to ensure all incoming axons were visible. Images are shown as maximum intensity projections. NMJs were quantified as monoinnervated when only a single axon was observed, and as poly-innervated when 2 or more axons connected to the endplate. For perforation analysis, gaps in α-BTX staining within the endplate were quantified (Fig. 2E). All NMJ quantifications were only performed on ‘en-face’ endplates to avoid misquantification due to the imaging angle. All analyses were performed blind to the genetic status of the material. Statistical analysis was conducted in GraphPad Prism 6. We performed unpaired, two-tailed t-tests and *P*-values were corrected for multiple comparisons with the Holm-Sidak method, with the exception of the analysis of extraocular muscles in Fig. 2D, where only one time point was investigated. Multiple testing correction for poly-innervation status was performed across time points, whereas for perforation studies, correction was applied across perforation status within each time point.

### Data access

All RNA-seq data generated in this study have been deposited at the Gene Expression Omnibus (GEO) of the National Center for Biotechnology Information, with the accession number GSE115706. Some samples utilized in this study (control SC P5 samples) were previously deposited into GEO with the accession number GSE76514. Samples that contain two raw data (.fastq) files were pooled during the mapping procedure.

## Acknowledgements

We would like to thank Mattias Karlén for his excellent work in creating the schematics in figures 1 and 7. We would also like to thank Marta Paterlini for providing excellent technical expertise regarding the Leica laser capture microscope. This work was supported by grants from the Swedish Research Council (2016-02112) to E.H., EU Joint Programme for Neurodegenerative Disease (JPND) (5292014-7500) to E.H. and R.S., Karolinska Institutet to E.H. C.S. is supported by a postdoctoral fellowship from the Swiss National Science Foundation.

## Author contributions

E.H. conceived the study, and E.H and R.S. supervised the project. S.N., H.S., R.S. and E.H. designed experiments. S.N., J.N., L.H.C., I.A., M.v.d.L, C.S. and Q.D. acquired data. S.N., H.S., J.N., C.S., R.S., and E.H. analyzed data. S.N. and E.H. wrote the manuscript with the help of H.S., J.N. and R.S. All authors edited and gave critical input on the manuscript.

## Disclosure declaration

The authors declare no conflicts of interest.

